# Target configuration determines how and what we learn during sensorimotor adaptation

**DOI:** 10.1101/2025.06.10.658797

**Authors:** Pamela Villavicencio, Jonathan S. Tsay, Cristina de la Malla

## Abstract

Motor adaptation—the process of correcting movement errors through feedback and practice—is a fundamental human capacity that keeps our actions well-calibrated amid changes in the environment and the body. However, how training context—specifically, the configuration of targets in the workspace—shapes how we learn and what we learn during motor adaptation remains unknown. To investigate this, we conducted two reaching experiments in which participants experienced a visuomotor gain perturbation, with feedback scaled to 1.3x (Exp 1) or 0.7x (Exp 2) the actual movement distance. In both experiments, participants were assigned to either the Extent Group, which trained with targets of varying amplitudes in a single direction, or the Angular Group, which trained with targets of equal amplitude in different directions. We found marked differences in *how* the two groups learned: the Angular Group learned more implicitly than the Extent Group, as evidenced by larger post-perturbation aftereffects when participants were instructed to forgo re-aiming strategies. Just as striking were the differences in *what* the two groups learned: the Angular Group learned a translation rule, which generalized to new directions but not amplitudes, while the Extent Group learned the imposed gain rule, which generalized to new amplitudes but not directions. Together, these findings underscore the importance of training context in determining how and what we learn.

## Introduction

Motor adaptation—the process of correcting movement errors through feedback and practice—is a fundamental human capacity that keeps our actions well-calibrated amid changes in the environment and the body (Shadmehr et al., 2010; Tsay et al., 2024). For example, when switching to a new computer mouse with a different gain sensitivity, the same hand movements may produce larger or smaller cursor displacements. Yet, within a short amount of time, we rapidly adapt to restore precise control over the cursor.

Motor adaptation is known to rely on multiple learning processes. Explicit strategies enable deliberate, flexible motor corrections, while implicit learning keeps the motor system finely tuned through unconscious, automatic adjustments (H. E. Kim et al., 2021; Maresch et al., 2021; Mazzoni & Krakauer, 2006; McDougle et al., 2016; Morehead et al., 2017; Morehead & Xivry, 2021; Taylor et al., 2014; Tsay et al., 2020, 2022). A growing body of research has characterized the distinct properties of implicit and explicit learning. For example, explicit strategies are flexible, scaling with different perturbation sizes. By contrast, implicit learning is rigid and largely insensitive to the size of the perturbation. Moreover, explicit strategies generalize broadly to new contexts, such as different target locations (McDougle et al., 2017; McDougle C Taylor, 2019). Implicit learning is again markedly rigid, generalizing only narrowly around the trained target (Day et al., 2016; Forano et al., 2021; Krakauer et al., 2000; Morehead et al., 2017; Wang & Taylor, 2021).

Despite these well-characterized properties, it remains unknown how training context—specifically, the configuration of targets—influences the balance between implicit and explicit processes. Standard visuomotor adaptation tasks have revealed that target configuration can bias performance (Taylor & Ivry, 2013). For example, grid-like arrangements tend to promote translation-based rules, while circular layouts bias participants toward rotational rules—even when the underlying transformation is always rotational. However, previous studies have not separated the contributions of implicit and explicit learning. Thus, target layout may have no impact on the balance between learning processes, with implicit and explicit systems contributing similarly across configurations. Alternatively, specific target layouts may preferentially recruit one system over the other.

To tackle this, we conducted two reaching experiments in which participants experienced a visuomotor gain perturbation, with feedback scaled to 1.3x (Exp 1) or 0.7x (Exp 2) the actual movement distance. In both experiments, participants were assigned to either the Extent Group, which trained with targets of varying amplitudes in a single direction, or the Angular Group, which trained with targets of equal amplitude in different directions. Both groups trained on a shared target, enabling us to cleanly compare how different training contexts influenced the explicit and implicit components of adaptation. Additionally, we included no-feedback reaches to novel targets, instructing participants to either engage or withhold their explicit re-aiming strategies; this approach allows us to isolate implicit and explicit contributions to context-dependent learning, revealing which system supports generalization across novel extents and directions—and probing whether participants learned a gain-based or an alternative representation. Together, this work deepens our understanding into how training context—specifically, the configuration of targets—determines not only *how* we learn, but also *what* we learn during motor adaptation.

## Methods

### Participants

42 unique participants took part in Experiment 1 (mean age = 21 years) and 47 in Experiment 2 (mean age = 26 years). All participants had normal or corrected-to-normal vision, no known motor impairments, and were predominantly right-handed, as assessed by the Edinburgh Handedness Inventory (Oldfield, 1971). Participants signed an informed consent prior to the start of the experiment and were compensated with course credit. Each participant was randomly assigned to one of two groups (Extent or Angular, see Experimental Design). This study was part of an ongoing research project approved by the Ethics Committee of the University of Barcelona (IRB: 00003099).

### Apparatus

Participants performed the task on a graphic tablet (Calcomp Drawing Board III 24240, 61 × 46 cm), that recorded the position of a hand-held stylus at 145 Hz. Stimuli were projected at a frame rate of 75 Hz and a resolution of 1024 × 768 pixels onto a back-projection screen. A half-silvered mirror placed between the screen and the tablet reflected the stimuli while occluding the participant’s view of their hand. This created the perception that both the hand and stimuli occupied the same plane on the tablet surface. LED lights underneath the mirror modulated the visibility of the arm and were off to occlude vision of the arm. The lights in the room always remained off. A computer controlled the presentation of the stimuli and registered the position of the stylus. The experiment was programmed using custom code in Python.

### General Procedure

Participants performed quick center-out reaching movements from a gray home position (1 cm diameter) to several possible blue targets (1 cm diameter). To begin each trial, participants placed a white cursor (0.6 cm diameter), which was aligned with the stylus, on the home position located at the center of the tablet. After remaining on the home position for a random interval between 0.8 and 1 second, the home position disappeared, and a target appeared. Participants were instructed to make a quick, ballistic reaching movement to the target. The target was displayed for only 1 second, requiring participants to complete their reach before it disappeared. Following a 0.3-second delay after the target vanished, the home position reappeared, signaling participants to return their hand to the center of the workspace. The veridical cursor was provided when participants were within 1 cm of the home position, allowing them to reposition the stylus and initiate the next trial.

To minimize any impact on explicit learning, we imposed no constraints on reaction time—defined as the interval from target onset to movement initiation (when hand velocity exceeded 15% of peak velocity) (Maresch et al., 2021). However, to discourage online corrections, participants saw a “too slow” message for 0.75 seconds if movement time—defined as the interval from movement initiation to termination (when velocity dropped below 10% of peak)—exceeded 0.5 seconds.

Feedback was presented in one of three forms: veridical, perturbed, or absent. The cursor was aligned with the stylus with veridical feedback. When the feedback was perturbed, it was scaled by a gain of either 1.3x (Exp 1) or 0.7x (Exp 2), requiring participants to shorten or extend their reach to hit the target, respectively. When the feedback was absent, participants did not receive any visual feedback during their reach.

### Experimental Design

Participants in both experiments were assigned to one of two groups: Extent or Angular (Fig. 1A). The Extent group trained with two targets of differing amplitudes (7 and 11 cm) that were either along the cardinal (90º) or diagonal (315º) axis (counterbalanced across participants accounting for reach biases) (Wang et al., 2024). The Angular group trained with two targets in different directions (counterbalanced across participants, cardinal: 90º and 180º; diagonal: 45º and 315º), but with the same amplitude (11 cm). For reference, 0º is defined as the direction pointing to the right from the home position.

**Figure 1.**
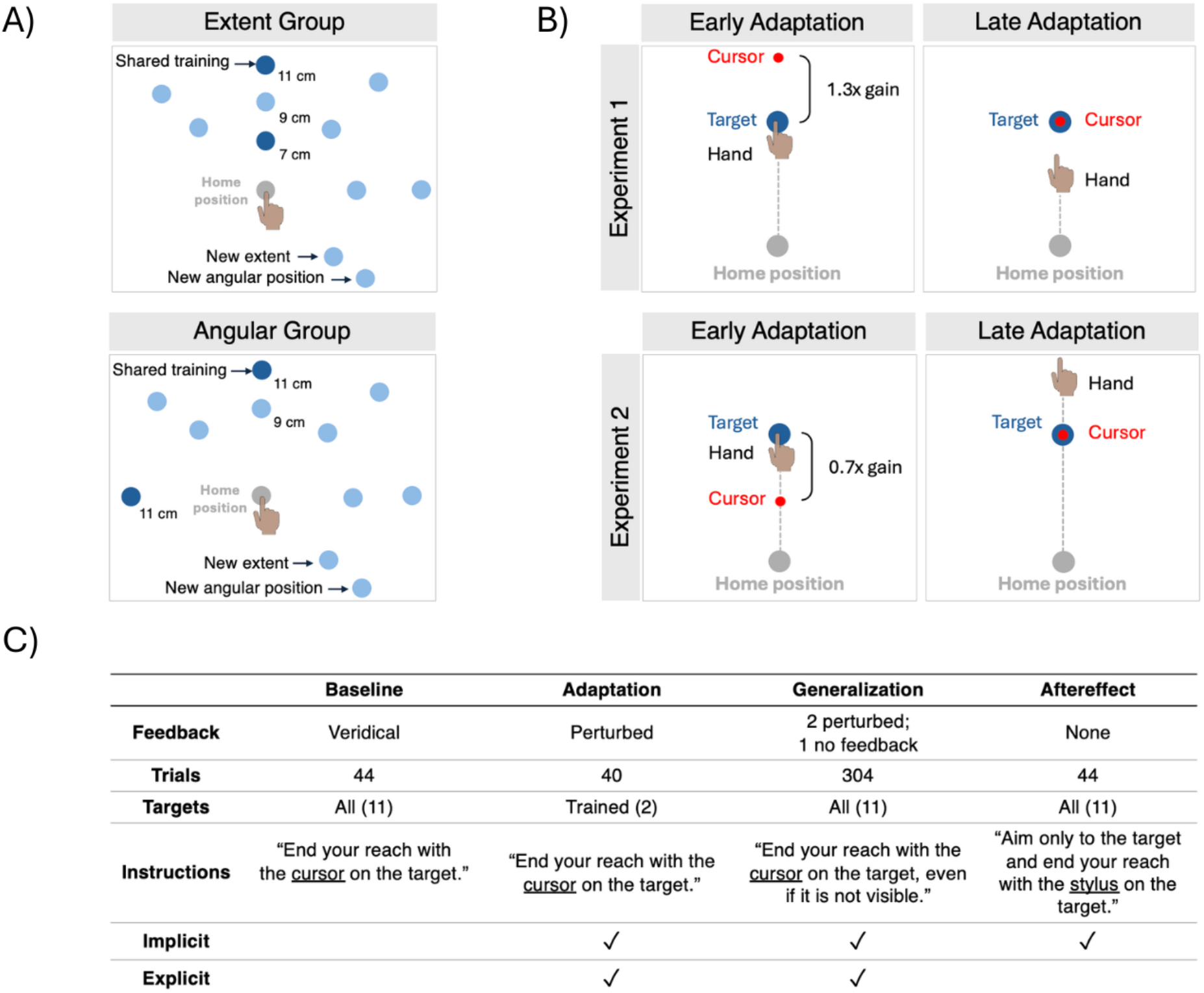
Experimental Design. **(A)** Target configurations for the Extent and Angular Group. Each group had two training targets (dark blue). The Extent group trained on two targets of differing amplitudes (7 and 11 cm) that were either along the cardinal (90º) or diagonal (315º) axis (counterbalanced across participants). The Angular group trained with two targets in different directions (counterbalanced across participants, cardinal: 90º and 180º; diagonal: 45º and 315º), but with the same amplitude (11 cm). For reference, 0º is defined as the direction pointing to the right from the home position. The 10 generalization targets (light blue)—including the shared trained target—were identical across groups. The only difference between groups was the unshared training target. Generalization targets were 9 and 11 cm from the home position. In the actual experiment, all targets appeared the same shade of blue. **(B)** Adaptation Schematic. Under a visuomotor gain perturbation, participants initially see the cursor either overshoot (Exp 1) or undershoot (Exp 2) the target (left panel: early adaptation). To compensate and bring the cursor onto the target, they must shorten their reach in Exp 1 or extend it in Exp 2 (right panel: late adaptation). **(C)** Experimental blocks. In the baseline block, participants reached to both training and generalization targets with veridical cursor feedback. During the adaptation block, participants reached to only the training targets, receiving perturbed visual feedback (Exp 1: 0.7x gain; Exp 2: 1.3x gain). In the generalization block, participants received perturbed feedback at the training targets and no feedback at the generalization targets. Finally, the aftereffect block consisted of no-feedback reaches to all targets.

Both groups trained on a shared 11 cm target location and were tested on the same set of 10 generalization targets, including the trained target. Generalization targets were located at angular distances of 0º, 45º, 90º, 135º relative to the shared training target, at amplitudes of either 9 cm or 11 cm (Fig. 1A).

Both experiments consisted of four blocks (Fig. 1C: 432 total trials; 25 minutes): baseline, adaptation, generalization, and aftereffect. Baseline consisted of 44 trials with veridical feedback to the generalization and training targets (11 total targets). Adaptation consisted of 40 trials with perturbed feedback (Exp 1: 1.3x gain; Exp 2: 0.7x gain) to the two training targets. The generalization block consisted of 304 trials, including 204 perturbed-feedback trials to the training targets and 100 no-feedback trials to the generalization targets. Specifically, each of the 10 generalization targets was presented 10 times in random order, with each trial preceded by two perturbed-feedback trials to the training target. Aftereffect consisted of 44 trials with no feedback to all targets (11 total targets). Instructions preceding each block are detailed in Figure 1C.

### Data Analysis

All data and statistical analyses were performed in R. The primary dependent variable in our analyses was the reach amplitude, defined as the straight-line distance from the home position to the stylus position at the end of the reach. We baseline-corrected each participant’s reach amplitude by subtracting the mean amplitude error from the second half of baseline trials for each target. Amplitude error was defined as the difference between the participant’s reach amplitude and the target amplitude. Euclidean error was defined as the distance between the target position and the stylus endpoint at movement termination.

We excluded trials under the following conditions: (1) the stylus lost contact with the tablet (Exp 1: 0.45%; Exp 2: 0.82%); (2) the start and/or end of the movement was not detected (Exp 1: 0.45%; Exp 2: 1.19%); or (3) the reach amplitude or Euclidean error deviated by more than 3 standard deviations from a 5-trial moving average (Exp 1: 0.14%; Exp 2: 0.19%). In total, we removed 1.05% of trials in Experiment 1 (0% – 3.94% per participant). In Experiment 2, two participants were excluded for having more than 20% of trials removed in at least one block. This yielded a final sample of 45 participants (Extent group: N = 23; Angular group: N = 22), with a total of 1.47% of trials excluded from these participants (0%–6.7% of trials removed per participant).

Performance was summarized across four learning phases: baseline, late adaptation, late generalization, and aftereffect. Baseline included the second half of trials to each target in the baseline block (22 trials). Late adaptation comprised the last 10 trials to each training target in the adaptation block (20 trials). Late generalization included the second half of the generalization block: the last 102 perturbed-feedback trials to the training targets and the last 5 no-feedback trials to each generalization target. The aftereffect phase included all trials from the aftereffect block (44 trials).

We also quantified combined, implicit, and explicit learning. Combined learning was reflected in performance during late generalization with perturbed feedback. Explicit learning was estimated as the difference between late generalization and aftereffect performance, while implicit learning was defined as the remaining difference between aftereffect and baseline.

To examine how training context influenced learning at the trained (shared) location, we conducted a linear mixed-effects model (R: lmer function) with learning phase (baseline, late adaptation, late generalization perturbed-feedback, aftereffect) and group (Extent, Angular) as fixed (interacting) factors, participant ID as a random factor, and mean reach amplitude as the dependent variable. We also assessed the impact of training context on generalization to novel probe locations using a linear mixed-effects model with learning phase (baseline, late generalization no-feedback, aftereffect), group, angular distance from the trained direction (0º, 45º, 90º, 135º), and target amplitude (9, 11 cm) as fixed factors, and participant ID as a random factor. To test for learned representations at a novel extent in the trained direction, we fit a mixed model with solution (observed, gain, translation, location) and group as fixed factors and participant ID as a random factor.

We tested for overall effects using the anova function in R and conducted post-hoc two-tailed t-tests using the *emmeans* function with Tukey corrections. Effect sizes are reported as *η*^2^ for fixed factors, Cohen’s *D*_*z*_ for within-subjects t-tests, and Cohen’s *D* for between-subjects t-tests.

## Results

### Experiment 1: Extent training led to less implicit adaptation and greater reliance on explicit re-aiming compared to Angular training

We asked whether the spatial layout of training targets influences the relative engagement of implicit and explicit learning processes. Two groups of participants were exposed to the same visuomotor gain perturbation, in which the cursor was scaled to 1.3x the amplitude of their hand movements. In the Extent group, participants trained with two targets along the same direction but at different amplitudes. In the Angular group, participants trained with two targets at the same amplitude but in different directions. Both groups shared a common training target located 11 cm from the home position, allowing us to compare adaptation at an identical location between groups. To adapt successfully, participants must scale down their movements to land the perturbed cursor on the target.

At the shared target, both groups rapidly adapted to the gain perturbation, reducing their movement amplitude to bring the perturbed cursor on target (Fig. 2A) (main effect of phase: *F*(3,120)=134.89, *p* < 0.0001, *η*^2^ = 0.77; baseline vs late adaptation: *t*(120)=17.70, *p* < 0.0001, *D*_*z*_ *=* 3.86). Neither group fully adapted, likely due to the challenge of executing low-amplitude movements precisely (Albert et al., 2021; Borish et al., 2020).

**Figure 2.**
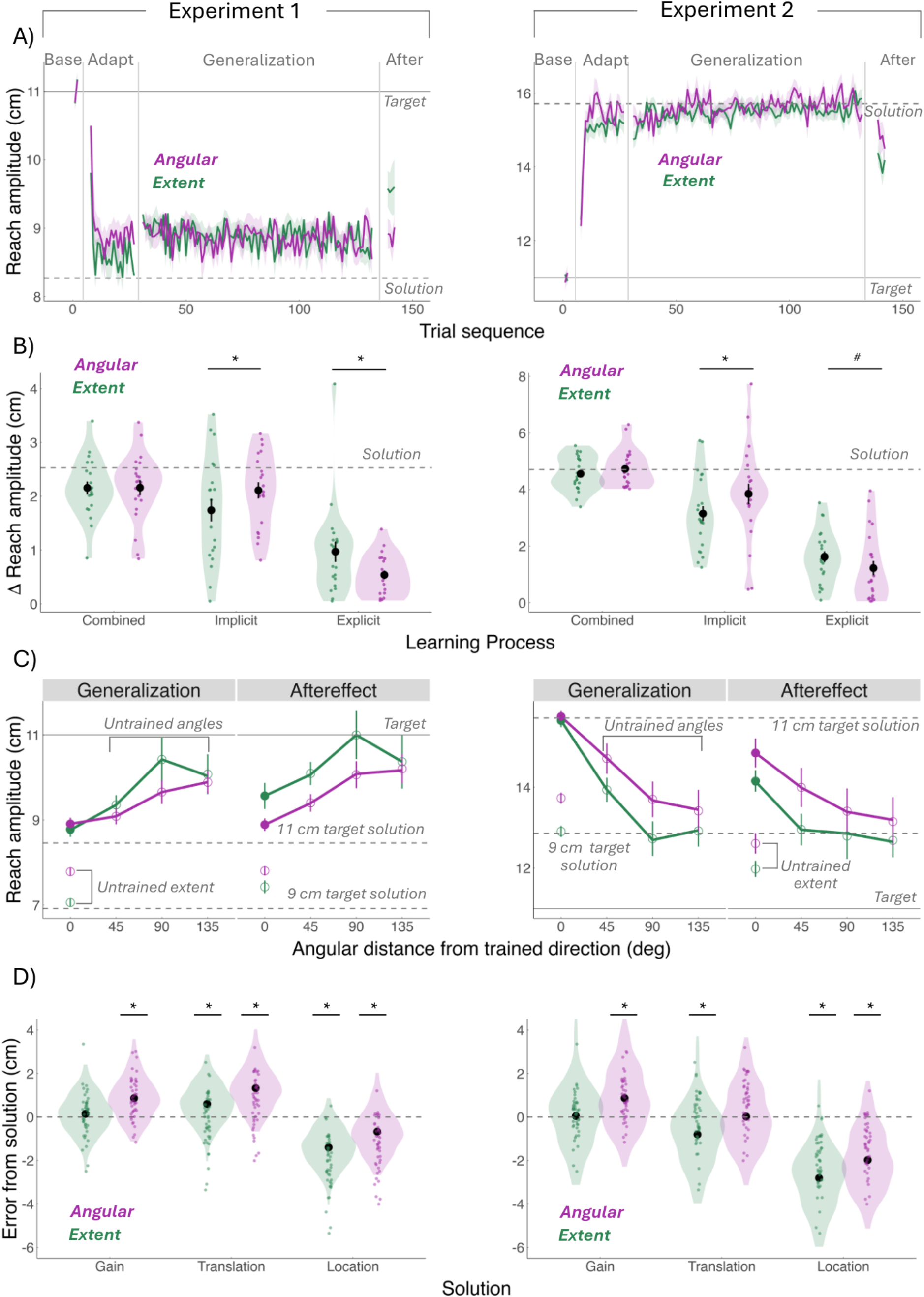
Extent training led to less implicit adaptation and greater reliance on explicit re-aiming compared to Angular training. **(A)** Mean reach amplitude time courses at the shared training target for the Angular (purple) and Extent (green) groups. Solid horizontal lines indicate the actual target amplitude (11 cm); dashed lines indicate the required amplitude to counteract the gain perturbation (Exp 1: 8.46 cm; Exp 2: 15.71 cm). Shaded error bars denote standard error. **(B)** Decomposition of learning at the shared training target. Total learning was quantified as the absolute change in reach amplitude during late generalization relative to baseline. Implicit learning was quantified as the difference between aftereffect and baseline, and explicit learning as the difference between late generalization and aftereffect. Dashed lines indicate the expected reach amplitude change for full adaptation to the gain perturbation. Translucent dots represent individual means; black dots indicate group means ± standard error. Asterisks (*) denote a significant difference between groups (*p* < 0.05); pound signs (#) indicate trending differences (p < 0.15). **(C)** Generalization to untrained directions and amplitudes. Mean reach amplitudes during late generalization no-feedback and aftereffect phases. Filled circles indicate the shared trained target; open circles represent novel targets. Dashed lines show the amplitudes required for full compensation at 9 cm and 11 cm targets, and the sold line indicates the actual 11 cm target amplitude. **(D)** Error relative to expected solutions. The difference between participants’ reach amplitude and the amplitude required for each solution. Negative values indicate undershooting; positive values indicate overshooting. Asterisks (*) denote a significant difference from the expected solution (p < 0.05).

Adaptation was sustained throughout the generalization phase (baseline vs late generalization: *t*(120)=16.77, *p* < 0.0001, *D*_*z*_ *=* 3.66). To isolate implicit learning, participants were instructed to stop re-aiming and simply reach to the original 11 cm target. Both groups showed robust aftereffects, indicating an implicit contribution to learning (baseline vs aftereffect: *t*(120)=13.77, *p* < 0.0001, *D*_*z*_ *=* 3.01). Additionally, reach amplitude increased from late generalization to aftereffect (*t*(120)= -3.00, *p* = 0.017, *D*_*z*_ = -0.65), suggesting that explicit re-aiming also contributed to adaptation. Notably, implicit learning accounted for the majority of total adaptation in both groups: 80.93% in the Extent Group and 97.69% in the Angular Group.

Given that both training groups engaged implicit and explicit learning processes, we turned to our central question: Does training context differentially engage these processes? We observed a marked interaction between Group and Phase (*F*(3,120)=5.36, p = 0.0017, *η*^2^ = 0.12). First, both groups exhibited similar levels of adaptation (Extent vs Angular, late adaptation: *t*(119)= -1.46, *p*= 0.148, *D*= -0.55; late generalization: *t*(119)= 0.02, *p*= 0.98, *D*= 0.009), indicating that both training schedules were equally effective in driving learning performance.

Second, however, the Extent group exhibited reduced implicit aftereffects compared to the Angular group (Extent vs Angular, aftereffect: *t*(119)= 3.02, *p*= 0.003, *D*= 1.15). Correspondingly, the Extent group showed a marked drop from late generalization to aftereffect (*t*(120)= -3.96, *p*= 0.0031, *D*_*z*_ = -1.22)—a hallmark of explicit strategy use— whereas this drop was absent in the Angular Group (*t*(120)= -0.28, *p* = 1.00, *D*_*z*_ = -0.09) (Fig. 2B). Indeed, there was a significant between-group differences in this drop in performance between late generalization and aftereffect phases (Extent vs Angular: *t*(120)= 2.61, *p* = 0.0103, *D* = -0.74), underscoring how the Extent group relied more on explicit re-aiming compared to the Angular group.

Together, while both groups learned primarily in an implicit manner, Extent training elicited relatively less implicit adaptation and greater reliance on explicit re-aiming compared to Angular training. In short, the spatial layout of training targets determined *how* people learned.

We then asked whether training context influenced *what* people learned—that is, how the participants represented the perturbation. To test this, we examined how learning generalized to novel target locations that differed in either extent (9 cm) or angular direction (0º, 45°, 90º, 135º) from the shared training target.

We first examined generalization to a novel extent (9 cm) while keeping the movement direction constant (0º). The Extent group fully generalized, accurately scaling their movements to the new target (Fig. 2C), whereas the Angular group exhibited partial generalization, overshooting by 0.9 cm (*t*(223)=-1.88, *p*= 0.06, *D* = -0.74). Strikingly, this pattern reversed when generalizing to novel angular directions (45°, 90º, 135º) with a constant extent (11 cm): the Angular group generalized more effectively than the Extent group (Fig. 2C; *t*(69.4)=0.95, *p*= 0.34, *D* = -0.27). Together, these results underscore the principle of training-specific generalization—Extent training promotes transfer to new amplitudes, while Angular training supports transfer to new directions.

But did both groups learn the same perturbation—or did they form different representations of it? If the 1.3x gain was represented veridically—as a scaled mapping— then a movement toward a 9 cm target should result in a reach of 6.9 cm. If the perturbation was interpreted as a fixed, additive offset (translation), we would expect movements to land at 6.5 cm. Alternatively, if the perturbation was encoded as a discrete solution tied to the original 11 cm training target (location), movements should generalize to 8.5 cm.

Strikingly, the two groups represented the perturbation differently, as indicated by a significant interaction between group and solution (*F*(3,131)=9.55, *p*= 0.0000095, *η*^2^ = 0.18). The Extent group seemed to represent the perturbation as a gain: their reaches closely matched the scaled solution with no significant difference from the predicted gain-based movement (*t*(120) = 1.11, *p*=0.95, *D*_*z*_= 0.34), but significant deviations from the alternatives (translation: *t*(120)= 4.94, *p*= 0.0001, *D*_*z*_= 1.53; location: *t*(120)= -11.73, *p*< 0.0001, *D*_*z*_= -3.62). In contrast, the Angular Group did not show clear evidence of a distinct representation, as their behavior significantly deviated from all predicted patterns (gain: *t*(120)= 7.39, *p*< 0.0001, *D*_*z*_ *=* 2.25; translation: *t*(120)= 11.12, *p*< 0.0001, *D*_*z*_ *=* 3.43; location: *t*(120)= -5.55, *p*< 0.0001, *D*_*z*_ *=* -1.71) (Fig. 2D). Indeed, the difficulty of executing small-amplitude movements, combined with the similarity between gain- and translation-based solutions, likely obscured what the Angular Group was learning— prompting us to design Experiment 2 to address these limitations.

In summary, training context – specifically the spatial layout of training targets – determines both how people learn and what they learn: Extent training relied more on explicit strategies and produced a gain-based representation, while Angular training engaged more implicit learning, though its underlying representation remains unclear. These results suggest that the spatial layout of targets not only affects how people adapt—but also fundamentally changes what is learned.

### Experiment 2: Extent training promotes gain-based learning, whereas Angular training favors translation-based representations

In our previous experiment, we were unable to determine how the Angular group represented the perturbation. We suspect this was due to two key factors: first, the representational solutions—particularly the translation-based and gain-based solutions—were closely spaced (i.e., reach amplitudes to 9 cm targets: translation = 6.46 cm; gain = 6.92 cm), making it difficult to distinguish between them; second, scaling down movements is inherently challenging, as small amplitudes are harder to execute precisely.

To address these limitations, we designed Experiment 2 using the same task structure as Experiment 1 but with a 0.7x gain (instead of 1.3x). This required participants to scale up their movements (instead of scale down). This inversion will not only improve the signal-to-noise ratio in motor execution but, crucially, will also increase the separation between the predicted outcomes of competing representational strategies (i.e., reach amplitudes to 9 cm targets: gain = 12.86 cm; translation = 13.71 cm; location = 15.71 cm), making it easier to determine how each group represented the perturbation.

We replicated all key learning patterns from Experiment 1. Specifically, both groups rapidly adapted to the gain perturbation by increasing their movement amplitude to bring the cursor to the target (Fig. 2A; main effect of phase: *F*(3,129) = 318.81, *p* < 0.0001, *η*^2^ = 0.88; baseline vs late adaptation: *t*(129)= -25.44, *p* < 0.0001, *D*_z_= -5.37). Interestingly, unlike in Experiment 1, both groups reached full adaptation—likely due to the larger, more easily controlled movements required by the 0.7x perturbation.

Moreover, both groups maintained adaptation through the late generalization phase (baseline vs. late generalization: *t*(129) = -27.58, *p* < 0.0001, *D*_*z*_ = -5.82), showed significant aftereffects (*t*(129) = -20.79, *p* < 0.0001, *D*_*z*_ = -4.38), and engaged explicit processes, as indicated by a significant drop from late generalization to aftereffect (*t*(129) = 6.80, *p* < 0.0001, *D*_*z*_ = 1.43). Importantly, both groups showed similar levels of total learning during late adaptation (Extent vs. Angular: *t*(156) = -1.11, *p* = 0.2671, *D* = - 0.50) and late generalization (*t*(156) = -0.66, *p* = 0.5091, *D* = -0.30), indicating that both training schedules were equally effective in driving successful performance.

Importantly, the groups differed in *how* they learned: the Extent group showed significantly smaller aftereffects than the Angular group (*t*(156) = -2.63, *p* = 0.0094, *D* = - 1.18), indicating reduced implicit learning. While both groups showed a significant drop in reach amplitude from late generalization to aftereffect (Fig. 2B; Extent: *t*(129) = 5.96, *p* < 0.0001, *D*_*z*_ = 1.76; Angular: *t*(129) = 3.68, *p* = 0.008, *D*_*z*_ = 1.11), the Extent group exhibited a trend toward a larger drop between phases (–1.40 cm) than the Angular group (–0.89 cm) (Extent vs Angular: *t*(129) = -1.54, *p* = 0.126, *D* = 0.38). Together, consistent with Experiment 1, Extent training resulted in less implicit adaptation and greater reliance on explicit strategies compared to Angular training.

We also replicated the key generalization patterns from Experiment 1: (1) Extent training produced greater generalization to novel amplitudes than Angular training (*t*(199)= -1.60, *p*= 0.11, *D*= -0.80); (2) Angular training yielded greater generalization to novel directions (*t*(68.6)= -1.55, *p*= 0.13, *D*= -0.58); and (3) as in training, generalization was primarily supported by implicit learning (baseline vs aftereffect, Extent: *t*(989)= - 11.45, *p*< 0.0001, *D*_z_= -1.69; Angular: *t*(989)= -14.86, *p*< 0.0001, *D*_z_= -2.24), accounting for 69.10% in the Extent group and 81.40% in the Angular group. These findings reinforce the principle of training-specific generalization—Extent training promotes transfer to new amplitudes, while Angular training supports transfer to new directions.

To assess how participants represented the perturbation, we compared their actual reach amplitudes to the predicted amplitudes for each hypothesized solution. In Experiment 2, the 0.7x gain shifted the predicted reach amplitude to 12.9 cm for a gain-based solution and 13.7 cm for a translation-based solution. The predicted amplitude for the location-based solution was 15.7 cm.

Consistent with Experiment 1, we found that each group differed in what they learned (interaction between group and solution: *F*(3,140.7)=4.95, *p*= 0.0027, *η*^2^ = 0.10). Again, the Extent group seemed to correctly represent the perturbation as a gain (*t*(129)= 0.41, *p*= 1.00, *D*_*z*_*=* 0.12), with significant deviations from translation (*t*(129)= -4.35, *p*= 0.0007, *D*_*z*_= -1.28) and location-based solutions (*t*(129)= -15.54, *p*< 0.0001, *D*_*z*_= -4.58). Strikingly, unlike in Experiment 1, the Angular Group’s representation of the perturbation clearly aligned with a translation (*t*(129)= 0.15, *p* = 1.00, *D*_*z*_= 0.05), deviating significantly from gain (*t*(129)= 4.81, *p* = 0.0001, *D*_*z*_= 1.45) and location-based solutions (*t*(129)= - 15.54, *p* < 0.0001, *D*_*z*_= -3.26). Together, these results demonstrate how the spatial layout of training targets can fundamentally influence the nature of what is learned.

## Discussion

We found that training context—specifically, the spatial layout of targets—shapes both how people learn and what they learn. This claim is supported by three key findings. First, training across different extents led to reduced implicit adaptation and greater reliance on explicit re-aiming, compared to training across different directions. Second, learning was context-specific: Extent training enhanced generalization to novel amplitudes, while Angular training enhanced generalization to novel directions. Third, Extent training promoted the imposed gain-based representation, whereas Angular training favored a translation-based representation.

Our results align with prior studies showing context-dependent generalization (Taylor & Ivry, 2013)—specifically, that extent training supports generalization to new amplitudes, while angular training supports generalization to new directions. Building on this, we extend this work in two important ways. First, previous work confounds target layout with required motor solution—for example, applying the same rotational perturbation to circular and grid-like target configurations results in different compensatory movements. This makes it difficult to determine whether generalization differences arise from the spatial layout, the motor solution, or both. To address this, we chose the gain-based perturbation so that the required motor solution was identical across training contexts: in both extent and angular training, the compensatory movement in the task-relevant dimension—extent—was matched. This allowed us to demonstrate that target configuration alone can drive context-dependent generalization.

Second, unlike previous studies that did not isolate learning processes, we provided instructions to clearly dissociate implicit from explicit learning. Specifically, before the aftereffect block, participants were instructed to stop using any re-aiming strategy and reach directly to the visual target. This approach allowed us to demonstrate that different target layouts differentially engage implicit and explicit processes— challenging the conventional assumption that these systems contribute uniformly across different contexts.

Why might training context bias the engagement of implicit and explicit learning processes? We have several hypotheses: First, spatial layout may specifically modulate the implicit learning system. For example, movements constrained by extent and angle may engage different populations of sensorimotor neurons that may vary in both their capacity and rate of plasticity (Caceres et al., 2024; Kantak et al., 2013). Our results suggest that neurons tuned to extent-based movements may adapt more slowly, leading to weaker implicit learning, whereas neurons engaged by angular-based movements adapt more rapidly, supporting greater implicit learning.

Second, spatial layout may specifically modulate explicit learning. An extent-based layout may naturally prime a gain-based rule—one that aligns with the amplitude-deviating structure of the perturbation—which participants can readily adopt to reduce error (Hiatt, 2017, 2023). If implicit and explicit systems compete to reduce motor errors (Albert et al., 2022; Miyamoto et al., 2020), the Extent group has less residual error available to drive implicit adaptation, thereby indirectly suppressing it. In contrast, an angular layout—being incongruent with the relevant dimension for gain perturbations— may not prime extent-related rules and may evoke misleading associations (e.g., rotations), making it harder to form an effective explicit strategy.

Lastly, spatial layout may modulate general cognitive resources, which in turn impacts explicit learning. Specifically, Angular training may require greater attentional resources, as participants must simultaneously plan movement extent to compensate for the perturbation *and* determine direction based on target layout. This dual demand, coupled with increased uncertainty in motor planning, may hinder the discovery of a successful explicit strategy, increasing reliance on implicit learning (Forano et al., 2021; Heald et al., 2018; Howard et al., 2013, 2015; S. Kim et al., 2015; Ogasa et al., 2024). In contrast, extent training isolates a single relevant dimension—movement amplitude— simplifying both strategy formation and movement execution (Koshland et al., 1999; van Beers et al., 2004). Future studies are needed to disentangle these possibilities—whether spatial layout selectively biases one learning system or independently influences both.

Why did the Extent group accurately represent the perturbation, while the Angular group defaulted to a translation-based interpretation? Building on findings from numerical cognition showing that addition is typically easier to compute than multiplication, we propose that in ambiguous training contexts, participants default to a simpler additive rule—translation—as their action-outcome hypothesis. In contrast, when the context provides clearer cues, as in Extent training, they may override this default and adopt a more complex but accurate multiplicative, gain-based rule (Perfors et al., 2011).

This account predicts a trade-off: reasoning through a more complex but generalizable gain rule should result in slower learning relative to reasoning through a simpler but less generalizable translation rule. Yet, we did not observe any differences in learning rates between training contexts. This may be due to the overall task being too easy, driving uniformly fast, near-ceiling performance. Future studies using more complex perturbation structures may better expose the costs of sensorimotor reasoning and reveal how training context biases participants toward specific policies (Tsay et al, 2024).

## Acknowledgements

This work was supported by grants PID2020-116400GA-I00, PID2023-150883NB-I00 and CNS2022-135808 funded by MCIN/AEI/10.13039/501100011033 and by the European Union NextGeneration EU/PRTR to CM. PV was supported by grant PRE2021-097890 funded by MICIU/AEI/10.13039/501100011033 and by the FSE+. JST was funded by Carnegie Mellon University Department of Psychology Seed Funding. The authors would like to thank Manel Moreno for his help in programming the experiment.

